# A single-cell transcriptomic atlas of the pigtail macaque lung

**DOI:** 10.64898/2026.11.05.764748

**Authors:** Taeyun Kim, John A. Cornelius, Vanshika Sindhu, Tyla Duell, Sidney Sun, Sophia C. Chima, Hazel Huang, Orlando Cervantes, Raj P. Kapur, Hong Zhao, Briana M. Del Rosario, Nehmat Bhutani, Emlyn Fettig, Maria Waldorf, Devesh Bajaj, Amanda Li, Andrew E. Vo, Amy Gretch, Gygeria Manuel, Jeff Munson, Alexandra Christodoulou, Melissa R. Berg, Chris English, Audrey Baldessari, Kristina M. Adams Waldorf

## Abstract

**Background:** The lungs undergo substantial physiological and immunological changes during respiratory illness to meet the host’s increased metabolic and oxygen demands. These responses require coordination among epithelial, endothelial, stromal, and immune cells throughout the respiratory zone, interstitium, and vasculature. Nonhuman primates (NHPs) are a valuable model for studying pulmonary biology as they have similar anatomic, physiologic, and immunologic correspondence to humans. Despite the widespread use of NHPs in pulmonary research, the cellular diversity and molecular characteristics of the pigtail macaque lung remain incompletely defined.

**Methods:** Parenchymal lung tissue from five adult female pigtail macaques (*Macaca nemestrina*) was dissociated into single cells and profiled using 10x Genomics 3’ single-cell RNA sequencing (scRNA-Seq). Next, ambient RNA was removed and quality control analyses performed. Samples were then integrated using scVI and scANVI. Cell types were labeled after unsupervised clustering and subclustering, followed by manual annotation using canonical marker genes. We compared lineages and cell types with the Level 4 Human Lung Cell Atlas (HLCA) using scArches reference mapping and with the Rhesus Immune Reference Atlas (RIRA) using Seurat label transfer.

**Results:** The atlas includes 61,323 cells representing 37 cell types in epithelial, mesenchymal, endothelial, lymphoid, and myeloid populations. Annotated populations included both major and rare lung cell types, including alveolar type I and II cells, club and multiciliated cells, aerocyte capillary and lymphatic endothelial cells, pre-terminal bronchiole secretory cells, CD4+ and CD8+ T cells, and several innate-like lymphocyte populations (gamma delta T cells, MAIT-like cells, innate-like CD8+ T cells, and group 3 innate lymphoid cells). There was strong concordance at HLCA annotation level 4, including AT1 and AT2 cells (96% and 93%), B cells (94 to 97%), CD4+ T cells (99 to 100%), classical and non-classical monocytes (84% and 85%), alveolar macrophages (80%), and mast cells (96%). Immune populations mapped to the RIRA with similar fidelity (alveolar macrophages and B cells, 99%).

**Conclusions:** While studies of pandemic respiratory infections, vaccines, and therapeutics rely on NHP disease models, this atlas provides a unique reference for epithelial, endothelial, stromal, and immune cell states in the pigtail macaque lung. We found that single-cell populations from the pigtail macaque lung are highly similar to those in the human lung at the lineage level. This reference will enable studies of pathogenesis and vaccine and drug efficacy in a highly translational model of the human lung.

## Background

The lung is a complex organ responsible for gas exchange, maintenance of tissue homeostasis, and immune surveillance.^1, 2^ During respiratory illness, the lung undergoes substantial physiological and immunological changes to meet the host’s increased metabolic and oxygen demands. These modifications require coordinated responses among epithelial, endothelial, stromal, and immune cells throughout the respiratory zone, interstitium, and vasculature.^3–5^ Experimental models are essential for understanding lung disease pathogenesis across cell types and for testing vaccines and therapeutics against pandemic respiratory viruses.^6^ Nonhuman primates (NHPs) are valuable for these studies because macaques closely resemble humans in lung anatomy, physiology, and immunology.^7–9^

The rhesus macaque (*Macaca mulatta*)^9, 10^, pigtail macaque (*Macaca nemestrina*), and cynomolgus macaques (*Macaca fascicularis*)^10, 11^ are among the most common models for investigating pulmonary physiology and respiratory infections. However, despite their widespread use in pulmonary research, the cellular diversity and molecular characteristics of NHP lungs remain incompletely defined with few single-cell lung atlases available.^12^ Further, genetic background, immune cell composition, and susceptibility to pulmonary infection differ among macaque species and among geographically distinct populations within a species.^13, 14^ Species-specific reference datasets are therefore needed to support rigorous comparisons with disease models, as well as the human lung.

Single-cell RNA sequencing (scRNA-seq) has enabled high-resolution characterization of cellular diversity and gene expression patterns, transforming the study of complex tissues like the lung. Single-cell atlases of the human lung have identified a variety of epithelial, endothelial, mesenchymal, and immune cell populations.^15, 16^ Progress has also been made in single-cell resources designed to characterize NHP lungs, including a cynomolgus lung atlas^12^, multi-tissue rhesus immune atlas containing lung cells^17^, developmental and preterm lung datasets^18, 19^, and infection- and disease-oriented datasets^20, 21^. However, existing datasets are often specific to a biological context. Immune-focused atlases often provide limited resolution of pulmonary epithelial, endothelial, and stromal compartments. Studies conducted on diseased lungs are designed to identify responses to specific experimental conditions. Moreover, most available NHP single-cell resources have focused on rhesus macaques or other NHP species, leaving the cellular composition and gene expression profiles of the normal pigtail macaque lung insufficiently characterized.

Our objective was to construct a scRNA-Seq atlas of the normal pigtail macaque lung. By characterizing endothelial, epithelial, mesenchymal, and immune cell populations at single-cell resolution, we provide a reference dataset for investigating cellular organization and transcriptional states in healthy pigtail macaque lung tissue. This reference dataset also provides a baseline comparison for future studies of pulmonary physiology, immunity, infection, injury, and disease. Our atlas complements immune-focused and disease-oriented NHP lung datasets with single-cell populations that align with human lung atlases.

## Results

### Comparative anatomy of the pigtail macaque and human lung

Annotating cell populations requires understanding the anatomical similarities and differences between the pigtail macaque and human lung (Fig. 1). The lung architecture between the two species is remarkably similar. Like the human lung, the pigtail macaque has conducting airways, terminal and respiratory bronchioles, alveolar ducts and sacs, and a mesothelial layer lining the pleural surface. These similarities led us to plan cell population labels that would align with the human lung.

**Figure 1.**
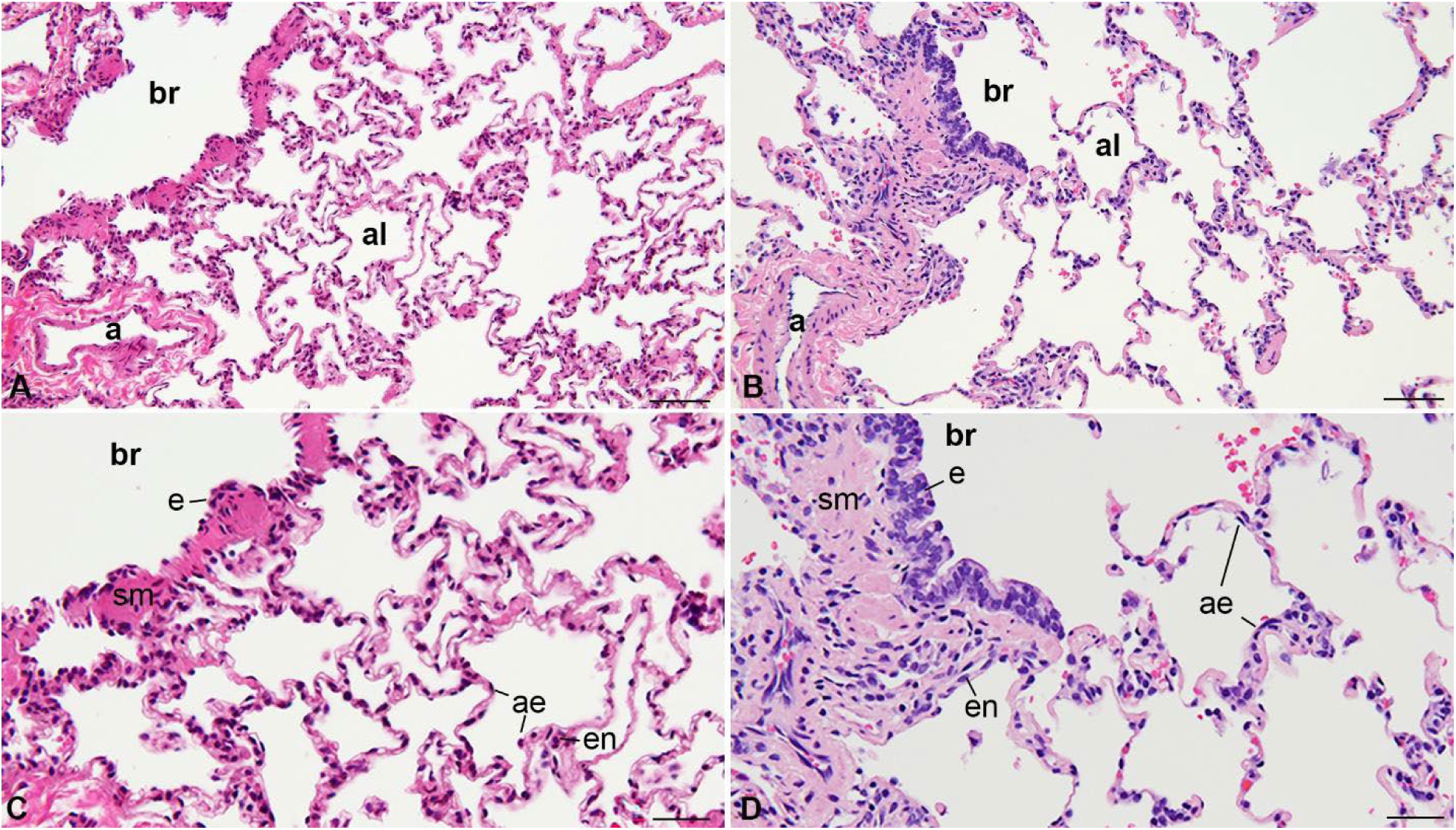
Comparative histopathology between the pigtail macaque and human lung Side-by-side photomicrographs of adult macaque (A, C) and human (B, D) lung showing terminal bronchioles (br), arteries (a), and alveoli (al). Cell types include bronchiolar epithelial cells (e), alveolar epithelium (ae), smooth muscle (sm), and endothelium (en). Scale bars: A-B, 50 µm; C-D, 25 µm.

### Study Design

To determine the single-cell transcriptional profiles of the pigtail macaque lung, we collected parenchymal lung tissues at necropsy from five adult females for single-cell dissociation and analysis using the 10x Genomics Chromium scRNA-Seq platform (Table S1). We processed samples through a quality control pipeline to remove ambient RNA, low-quality populations, and doublets. Then, samples were integrated using scVI and scANVI, followed by manual annotation using canonical markers.

### Clustering, Annotation, and Cell Origin

Before scVI and scANVI integration, cells were clustered in part by animal (Fig. S1A). After integration, cells were mixed within clusters across all five animals (Figs. S1B and S2). We removed clusters with low gene counts or with non-specific or ambient RNA signatures but retained neutrophils (NEUT) as they are an expected population in NHP models of pulmonary disease (Fig. S1C-E). The final atlas contained 61,323 cells across 37 cell types, as shown in Figure 2.

**Figure 2.**
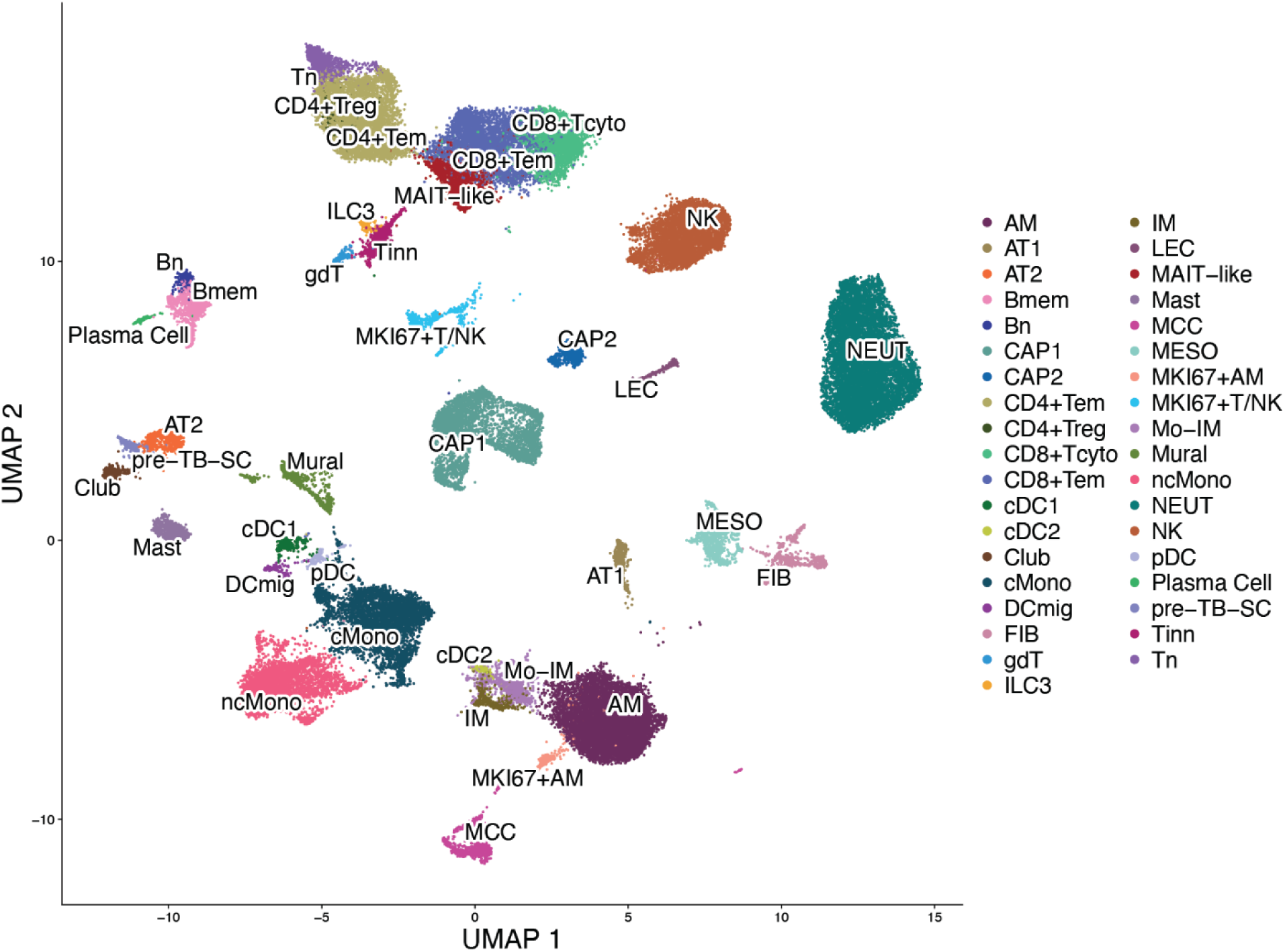
Single-cell atlas of the adult pigtail macaque lung

We resolved and annotated cell populations as closely as possible to the cell types in the Human Lung Cell Atlas (HLCA). Epithelial cells in the atlas included alveolar type 1 and 2 cells (AT1, AT2), Club cells (Club), multiciliated cells (MCC), and the pre-terminal bronchiole secretory cell (pre-TB-SC). Mesenchymal cells were fibroblasts (FIB), mural cells encompassing both pericytes and vascular smooth muscle cells (Mural), and mesothelial cells (MESO). We identified three endothelial cell populations: general capillary cells (CAP1), aerocyte capillary endothelial cells (CAP2), and lymphatic endothelial cells (LEC). There were 14 lymphocyte or innate lymphocyte populations: naïve T cells (Tn), proliferating T and natural killer cells (MKI67+T/NK), CD4+ T effector memory and regulatory T cells (CD4+Tem, CD4+Treg), CD8+ effector memory and cytotoxic T cells (CD8+Tem, CD8+Tcyto), innate-like T cells (Tinn), gamma delta (γδ) T cells, mucosal-associated invariant T-like cells (MAIT-like), group 3 innate lymphoid cells (ILC3), natural killer cells (NK), naïve and memory B cells (Bn, Bmem), and plasma cells (Plasma Cell). Finally, diverse myeloid populations were identified, including proliferating and non-proliferating alveolar macrophages (MKI67+AM and AM); interstitial macrophages (IM), monocyte-derived interstitial macrophages (Mo-IM), type 1 and type 2 conventional and plasmacytoid dendritic cells (cDC1, cDC2, pDC); classical monocytes (cMono); nonclassical monocytes (ncMono), migratory (mature) dendritic cells (DCmig), mast cells (Mast), and neutrophils (NEUT).

This UMAP shows clustering of single-cell transcriptomes, colored according to cell type, with each point representing an individual cell. Abbreviations: AM, alveolar macrophage; AT1, alveolar type 1 cell; AT2, alveolar type 2 cell; Bmem, memory B cell; Bn, naive B cell; CAP1, general capillary endothelial cell; CAP2, aerocyte capillary endothelial cell; CD4+Tem, CD4+ effector memory T cell; CD4+Treg, CD4+ regulatory T cell; CD8+Tcyto, CD8+ cytotoxic T cell; CD8+Tem, CD8+ effector memory T cell; cDC1, type 1 conventional dendritic cell; cDC2, type 2 conventional dendritic cell; Club, club cell; cMono, classical monocyte; DCmig, migratory dendritic cell; FIB, fibroblast; gdT, gamma delta (γδ) T cell; ILC3, group 3 innate lymphoid cell; IM, interstitial macrophage; LEC, lymphatic endothelial cell; MAIT-like, mucosal-associated invariant T-like cell; Mast, mast cell; MCC, multiciliated cell; MESO, mesothelial cell; MKI67+AM, proliferating alveolar macrophage; MKI67+T/NK, proliferating T and natural killer cells; Mo-IM, monocyte-derived interstitial macrophage; Mural, mural cell (pericyte and vascular smooth muscle cell); ncMono, nonclassical monocyte; NEUT, neutrophil; NK, natural killer cell; pDC, plasmacytoid dendritic cell; Plasma Cell, plasma cell; pre-TB-SC, pre-terminal bronchiole secretory cell; Tinn, innate-like T cell; Tn, naive T cell.

### Epithelial, Mesenchymal, and Endothelial Cells

We identified five epithelial cell populations in the parenchymal pigtail macaque lung tissue with elevated *EPCAM* (Fig. 3A–B). Both AT1 and AT2 cells were identified with active canonical markers similar to human lung atlases (Fig. 3B). Additionally, club cells expressed *SCGB1A1* and *MSLN,* and multiciliated cells (MCC) showed elevated *FOXJ1*, *RSPH1*, *DNAH5*, and *CAPS*. A fifth population expressed secretory genes (*SCGB3A2*, *SCGB1A1*) and AT2 genes (*SFTPC*, *NAPSA*, and *SFTPA1*), matching the known expression of pre-terminal bronchiole secretory cells (pre-TB-SC) in distal airways that give rise to AT2 cells.^22^ We did not identify goblet or basal cells, which was consistent with our sampling of parenchymal tissue.

**Figure 3.**
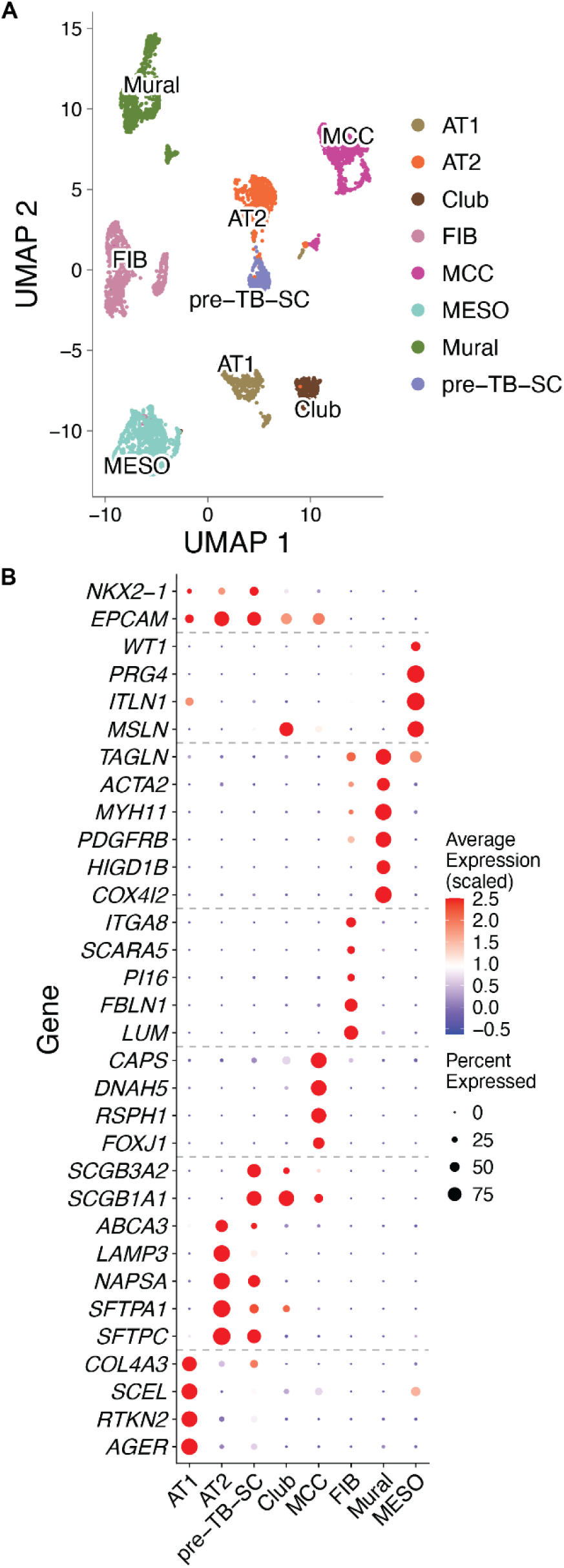
Epithelial and Mesenchymal Single Cells in the Pigtail Macaque Lung. Panel A shows UMAP visualizations of epithelial (AT1, AT2, pre-TB-SC, Club, MCC) and mesenchymal (FIB, Mural, MESO) single-cell populations. Each point represents an individual cell that is colored by identity. The dot plots show expression of canonical marker genes in these clusters (B), with dot size indicating the percentage of cells expressing the gene and color intensity scaled to average expression. Abbreviations are defined in the legend of Figure 2.

Three mesenchymal populations were identified. The first was a mixed fibroblast (FIB) population expressing *LUM* and *FBLN1* with markers of both adventitial (*PI16*, *SCARA5*) and alveolar (*ITGA8*) fibroblasts. The second was a mural cell population with active pericyte genes (*PDGFRB*, *HIGD1B*, *COX4I2*) and smooth muscle genes (*ACTA2*, *MYH11*, *TAGLN*). Third, we identified a mesothelial cell (MESO) population expressing *MSLN*, *ITLN1*, and *WT1*.

We observed three endothelial populations with elevated *PECAM1* and *CDH5* (Fig. 4A-B). General capillary endothelial cells (CAP1) had elevated *PTPRB*, *TEK*, and *EDN1,* with subsets showing transcriptional profiles of venous (*SELP*, *ACKR1*) and arterial (*HEY1*, *GJA5*) markers. Aerocyte capillary endothelial cells (CAP2) were also identified with expression of *EDNRB*, *HPGD*, *CYP2J2*, and *SULT1C4*. Finally, lymphatic endothelial cells (LEC) were annotated based on elevated *PROX1*, *CCL21*, *MMRN1*, *RELN*, and *LYVE1*.

**Figure 4.**
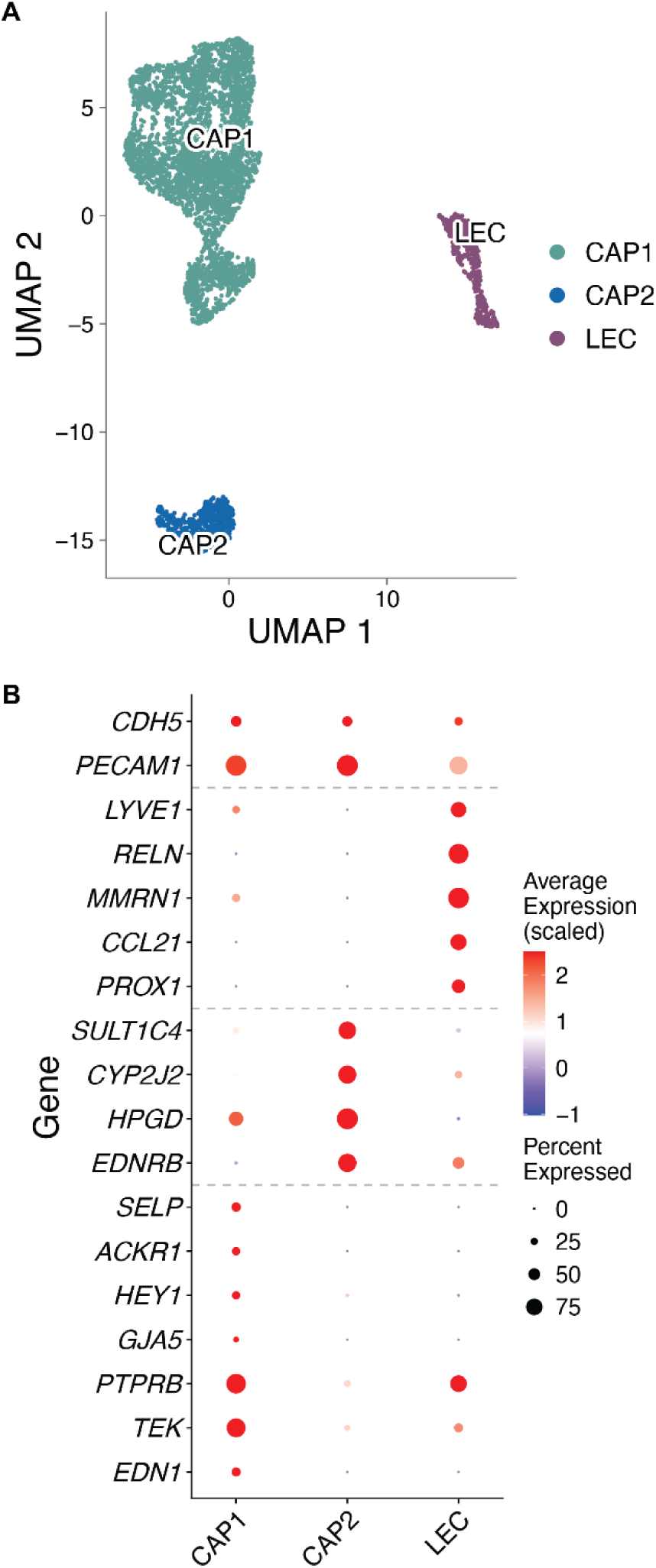
Endothelial Cells in the Pigtail Macaque Lung. Panel A shows a UMAP visualization of endothelial (CAP1, CAP2, LEC) single-cell populations. Each point represents an individual cell that is colored by identity. The dot plot in panel B shows expression of canonical marker genes in these clusters, with dot size indicating the percentage of cells expressing the gene and color intensity scaled to average expression. Abbreviations are defined in the legend of Figure 2.

### Lymphoid and Innate Lymphoid Cells

Defining lymphoid populations in a lung atlas is important to establish a reference for use in NHP models of pandemic respiratory infections. Eleven conventional and innate lymphoid single-cell populations were identified (Fig. 5A). The conventional T cells include naïve T cells (Tn), CD4+ effector memory T cells (CD4+Tem), CD4+ regulatory T cells (CD4+Treg), CD8+ effector memory T cells (CD8+Tem), and CD8+cytotoxic T cells (CD8+Tcyto). Innate and innate-like lymphocytes included natural killer (NK) cells, gamma delta T cells (γδT), mucosal-associated invariant T-like cells (MAIT-like), innate-like T cells (Tinn), and group 3 innate lymphoid cells (ILC3). The T cell populations expressed CD3E, and the CD4 and CD8 lineages expressed *CD4*, *CD8A*, and *CD8B,* similar to human T cells (Fig. 5B). All pigtail macaque T cells had elevated levels of CD69, which indicates residency in the lung.

**Figure 5.**
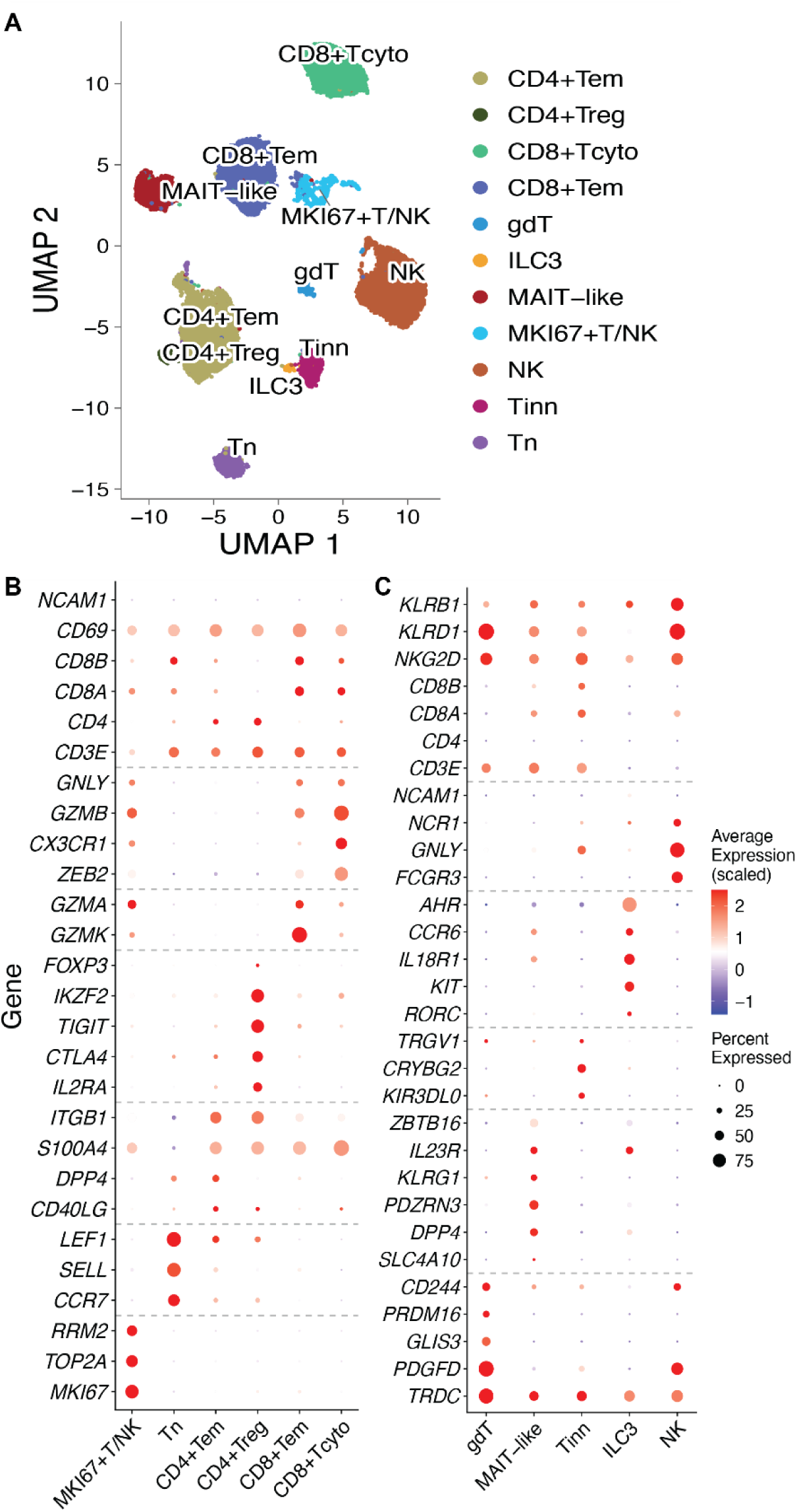
Lymphoid and Innate Lymphoid Single-Cell Populations in the Pigtail Macaque Lung Panel A shows a UMAP visualization of conventional and innate lymphoid populations (CD4+Tem, CD4+Treg, CD8+Tcyto, CD8+Tem, gdT, ILC3, MAIT-like, MKI67+T/NK, NK, Tinn, and Tn). Each point represents an individual cell that is colored by identity. The dot plots show expression of canonical marker genes in these clusters (B, C), with dot size indicating the percentage of cells expressing the gene and color intensity scaled to average expression. Abbreviations are defined in the legend of Figure 2. Note that we show *CD8B*, the annotation for ENSMMUG00000003532 in the Mmul10 reference (Table S2). *NKG2D* is also the annotation for the gene *KLRK1* in the Mmul10 reference.

Naïve T cells showed typical markers of quiescent T cells [*LEF1*, *SELL*(CD62L), *CCR7*] and lacked the gene signature of effector T cells (e.g., *GZMB*, *GZMK*; Fig. 5B). This cluster contained both CD4+ and CD8+ naïve T cells but could not be further resolved by unsupervised clustering. We also identified a non-cytotoxic effector, helper/memory CD4+Tem population with elevated *ITGB1* (CD29), *S100A4*, *DPP4* (CD26), and *CD40LG*, but lacking *GZMB*, *GZMK*, and *GZMA*expression. CD4+Treg cells were identified with canonical markers similar to those observed in humans [*IL2RA* (CD25), *TIGIT*, *CTLA4*, and *IKZF2* (Helios)]. The canonical marker FOXP3 was present in a small fraction of this population, consistent with poor RNA-level detection of Treg transcription factors in resting T cells.^17^ In the CD8+ effector populations, there were two discrete clusters that differed based on granzyme expression (Fig. 5B). CD8+Tem had elevated *GZMK* and *GZMA* but limited *GZMB* expression. In contrast, CD8+Tcyto strongly expressed *GZMB*, *CX3CR1*, and *ZEB2*, consistent with a terminally differentiated cytotoxic CD8+ T cell. The MKI67+T/NK population had elevated proliferation markers (*MKI67*, *TOP2A*), as well as *GZMA*, *GZMB,* and *CD8A*, indicating proliferating cytotoxic lymphocytes.

We annotated five innate lymphoid single-cell populations in the pigtail macaque lung (Fig. 5C). An NK cell population showed canonical markers of cytotoxic human NK cells [e.g., *FCGR3* (CD16), *KLRB1*, *KLRD1*, *KLRK1* (NKG2D), *GNLY*] and lacked *CD3E*. A group 3 innate lymphoid cell population (ILC3) was identified expressing *RORC*, *KIT* (CD117), *AHR*, *IL23R*, and *CCR6,* and lacking *CD3E* expression.

Three innate T cell populations showed a combination of *CD3E* and NK cell markers. A gamma delta T cell population (gdT) was identified co-expressing *CD3E* and *TRDC* with NK-associated receptors (*KLRD1*, *KLRK1*) and low *CD4*, *CD8A*, and *CD8B*. Other markers supporting this population included *CD244*, *PRDM16*, *GLIS3*, and *PDGFD*. We also annotated a population of innate-like CD8+ T cells (Tinn), which expressed *CD3E*, *CD8A*, and *CD8B, along with KLRD1 and KLRK1 (NKG2D, Fig. 5C).* This population is analogous to the EOMES+ KIR/NKG2A+ innate-like CD8+T cells described in human blood.^23^ Finally, we identified a mucosal-associated invariant T (MAIT)-like cell; in humans, MAIT cells represent a critical innate cell in the lung that responds rapidly to cells infected with pathogens that synthesize riboflavin (vitamin B2) metabolites. As the 3′ capture of poly A mRNA in our scRNA-Seq pipeline cannot resolve the TRAV1-2– TRAJ33/12/20 rearrangement that defines human MAIT cells, we designated this cluster as “MAIT-like” based on its transcriptional profile of *CD8A* (without *CD8B*) and *PDZRN3*, *KLRG1* (marker of terminal effector differentiation), and *DPP4* (regulates chemokine cleavage to modulate mucosal lymphocyte migration) expression.

Naïve B cells (Bn), memory B cells (Bmem), and plasma cells were also identified in the atlas (Fig. 6A). Universal B cell markers were present in both Bn and Bmem, including *MS4A1* (CD20), *CD79A*, *CD79B*, and *BANK1* (Fig. 6B). Bn was resolved by elevated *BACH2*, *CD72*, *FCER2*, *TCL1A*, and *IL4R*. Bmem was characterized by greater expression of *AIM2*, *TNFRSF13B* (TACI), *FCRL5,* and *SCIMP*. Plasma cells had a transcriptional profile consistent with antibody-secreting cells (*JCHAIN*, *MZB1*, *IGLL5*) and IgG heavy-chain constant genes (*IGHG1*, *IGHG3*).

**Figure 6.**
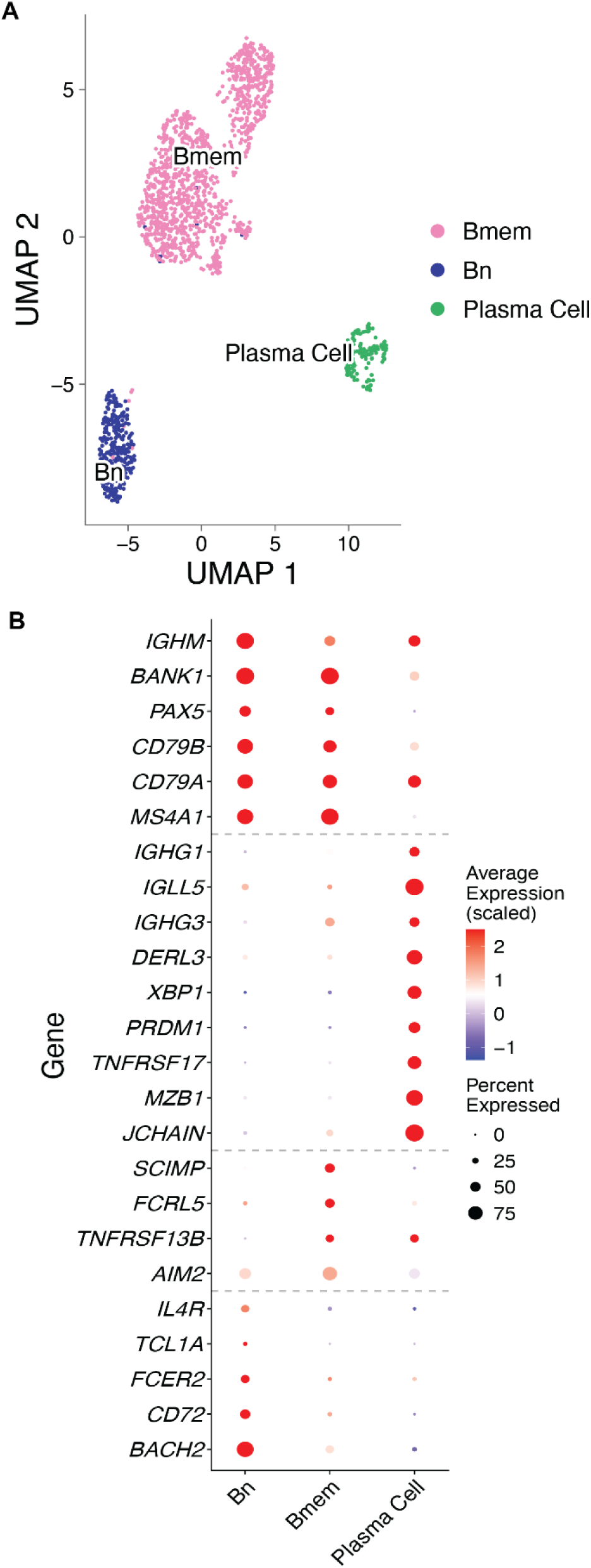
B Cell Populations in the Pigtail Macaque Lung Panel A shows a UMAP visualization of B cell populations (Bmem, Bn, and Plasma Cells) in the pigtail macaque lung. Each point represents an individual cell, colored by identity. The dot plots show expression of canonical marker genes in these clusters (B), with dot size indicating the percentage of cells expressing the gene and color intensity scaled to average expression. Abbreviations are defined in the legend of Figure 2.

### Myeloid Cells

We identified 12 myeloid populations that included two monocyte, four macrophage, four dendritic cell, one neutrophil, and one mast cell population (Fig. 7A). The classical monocyte (cMono) population expressed *CD14*, *VCAN*, and *S100A8/9*, while the non-classical monocyte (ncMono) expressed *FCGR3* (CD16) and *CX3CR1* (Fig. 7B). Macrophage populations shared expression of *CD14*, *CD68*, and *CD163* (Fig. 7B). Canonical markers for human alveolar macrophages (AM; *FABP4*, *PPARG*, and *MARCO*) and interstitial macrophages (IM; *C1QB* and *TMEM176B* but no *FABP4*) were also expressed in the pigtail macaque lung. A fourth population, monocyte-derived interstitial macrophages (Mo-IM), shared CD14, CD163 and CSF1R with IM and expressed SERPINB10, GPR183 and CLEC6A (Fig. 7B). Although the neutrophil population was low-quality, we retained it in the atlas as larger NEUT populations are anticipated in future datasets of respiratory infectious disease.

**Figure 7.**
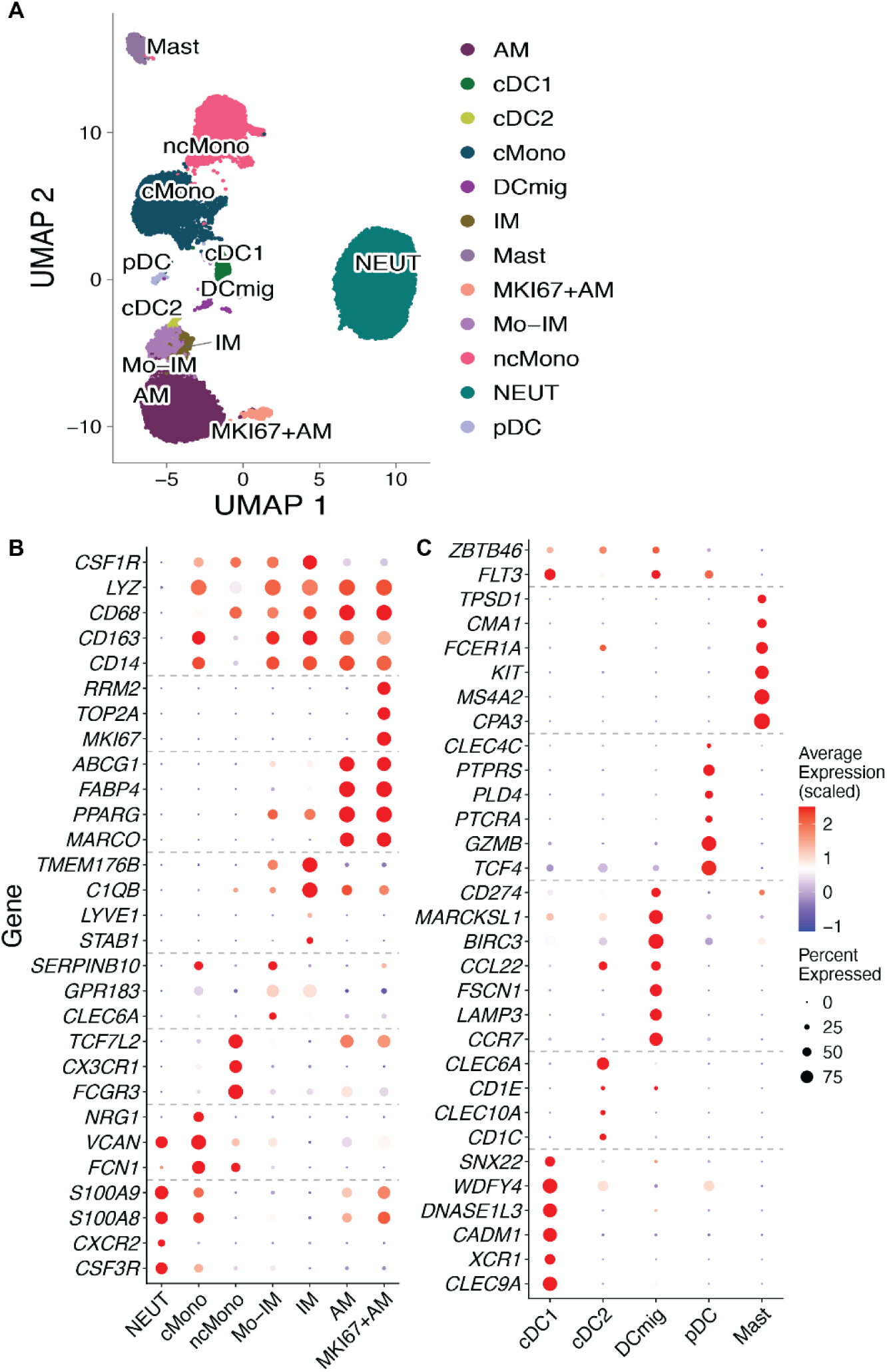
Myeloid Lineage. A UMAP visualization of myeloid populations (cDC1, cDC2, DCmig, pDC, AM, IM, Mo-IM, MKI67+AM, cMono, ncMono, NEUT, and Mast) is shown in panel A. Each point represents an individual cell that is colored by identity. The dot plots show the expression of canonical marker genes in these clusters (B, C) with dot size indicating the percentage of cells expressing the gene and color intensity scaled to average expression. Abbreviations are defined in the legend of Figure 2.

Four populations of dendritic cells were also resolved (Fig. 7C). A type 1 conventional dendritic cell cluster (cDC1) was identified, expressing *CLEC9A*, *XCR1*, *CADM1*, and *DNASE1L3* (Fig. 7C). Type 2 conventional dendritic cells (cDC2) expressed CD1C, CD1E, CLEC10A and CLEC6A (Fig. 7C). Mature migratory dendritic cells (DCmig) expressed *CCR7*, *LAMP3*, *FSCN1*, and *CCL22*. A fourth population of plasmacytoid dendritic cells (pDC) was characterized by *TCF4*, *GZMB*, *PTCRA*, and *CLEC4C* expression. Finally, a cluster of mast cells was found that expressed *CPA3*, *KIT*, *FCER1A*, and *CMA1*.

### Cross-Species Comparison with the Rhesus Macaque and Human Lung

To place the pigtail macaque lung in a cross-species context, we mapped its 37 cell types onto the core Human Lung Cell Atlas (HLCA) at Level 4, which was the level most comparable to the number and cell types in our atlas (Fig. S3).^15^ We also mapped the 26 immune cell types onto the Rhesus Immune Reference Atlas^17^ (RIRA, Fig. S4). When a reference lacked an equivalent population, cells were assigned to the closest available group. For example, neutrophils mapped to classical monocytes in the HLCA (100%), which has no neutrophil label, and to granulocyte monocyte progenitor-like cells in RIRA (83%).

Epithelial, endothelial, B cell and monocyte populations mapped to their expected counterparts with high fidelity. In the HLCA Level 4, AT1 and AT2 cells mapped to their human equivalents (96% and 93%), as did multiciliated cells (88%), CAP2 (86%) and mural cells (83%). B cells mapped to B cells in both references (94 to 99%). Classical and nonclassical monocytes mapped to their counterparts in RIRA (81% and 94%) and the HLCA Level 4 (84% and 85%). The AM population mapped to similar RIRA (99%) and HLCA (80%) populations.

Lower concordance reflected mainly differences in reference resolution rather than poor mapping. The HLCA Level 4 divides endothelial cells and fibroblasts into more subtypes than our atlas; the pigtail macaque CAP1 is split across three HLCA endothelial cell subtypes, while the FIB population mapped to three human fibroblast subtypes. RIRA assigned the pigtail IM (98%) and Mo-IM (81%) to alveolar macrophages, whereas the HLCA Level 4 assigned both mainly to interstitial macrophages (77% and 45%), with smaller fractions assigned to DC2 (IM, 18%; Mo-IM, 14%). Neither reference labels γδ T, MAIT or ILC3 cells separately. These mapped to HLCA Level 4 CD4 or CD8 T cells and to broad RIRA categories (“Gamma/Delta and Innate-like”: gdT and ILC3, 67% each; “Cytotoxic TCM-like”: MAIT-like, 89%). Our Tn population mapped in near-equal proportions to the CD4+ and CD8+ naïve T cells that RIRA separates (47% and 46%). Median mapping uncertainty was highest in innate-like T populations (gdT, ILC3 and Tinn) in RIRA and in cDC1, Mo-IM, IM and pre-TB-SC in the HLCA (Figs. S3-S4).

## Discussion

Diverse and rich single-cell populations were identified in the pigtail macaque lung with high similarity to the rhesus macaque and human lung. This atlas strengthens the value of the pigtail macaque as an NHP model of pandemic respiratory infections and lung disease. Resolution and careful annotation of cell types in this atlas makes it a useful reference for NHP studies of pandemic respiratory viruses (e.g., influenza A viruses, SARS-CoV-2) and inhaled toxins. Investigating how transcriptional states and proportions of single-cell populations change with infection and injury, which can provide valuable information regarding the pathogenesis and the preventive benefit of vaccines or therapeutic benefit of new drugs. While bulk RNA-Seq cannot resolve these differences to single-cell populations, and spatial profiling does not capture a larger volume of tissue with rarer cell types, single-cell resolution of cells in the pigtail macaque lung can provide a baseline for important studies in the context of disease, particularly pandemic respiratory infections.

Nonhuman primates (NHPs) are valuable models for respiratory disease and for evaluating vaccines and therapeutics against viral infections. Their lung anatomy, physiology, and mucosal immune responses resemble those of humans, and their size enables clinical imaging, repeated airway sampling, and pulmonary function testing alongside viral, immune, and pathological assessments.^6, 7, 24^ Rhesus macaques infected with SARS-CoV-2 developed transient respiratory disease, pulmonary infiltrates, and respiratory-tract viral replication, supporting studies of disease, vaccines, neutralizing antibodies as a correlate of protection, and remdesivir treatment.^25–29^ Pigtail macaque studies further showed variation in disease outcomes and immune signatures.^30^ For influenza research, cynomolgus macaque H1N1 studies informed challenge methods and vaccine evaluation^31^, while pigtail macaque H1N1 studies examined clinical signs, lung pathology, viral clearance, and host responses.^32–34^ H5N1 studies have linked clinical findings, viral measurements, lung pathology, and gene expression with disease severity, including severe lung injury in aerosol challenge models.^35–37^ A single-cell RNA-sequencing atlas of the pigtail macaque lung will provide a useful reference for future studies of respiratory viral infection and mucosal immunity in the context of nonhuman primate models of respiratory disease.

Studies in cynomolgus monkeys, African green monkeys, and rhesus and pigtail macaques have helped define host-pathogen interactions, pulmonary immune responses, and disease pathogenesis following SARS-CoV-2 infection.^25, 38^ These same models have also been used extensively to evaluate vaccine immunogenicity, protective efficacy, and antiviral therapeutics before advancement to human clinical trials for viruses such as influenza^39–43^ and respiratory syncytial virus^44–46^ (RSV). Macaque models are increasingly used to study many features of non-viral infections such as human pulmonary tuberculosis, including latent and active disease states, making them invaluable for studies of disease pathogenesis and countermeasure development.^47^ Recent advances in single-cell RNA sequencing have enabled high-resolution characterization of lung epithelial, stromal, and immune cell populations in healthy and diseased human lungs.^48, 49^ Similar advances have been made and are well developed for other species. However, despite these advances and the increased use of NHPs in translational and infectious disease research, comprehensive reference datasets remain limited for several NHP species, including the pigtail macaque. A comprehensive single-cell RNA-Seq atlas of the pigtail macaque lung will provide a valuable reference for studies of respiratory infection, inflammation, immune regulation, vaccine development, and therapeutic intervention.

Single-cell RNA-seq lung atlases are now extensive for humans and well developed for other species. Human atlases span adult, aging, fetal and newborn lung and have been integrated into reference resources of millions of cells.^15, 16, 50–53^ Travaglini et al. defined 58 cell populations in the human lung, and identified 17 molecular cell types that have been gained or lost during lung evolution in comparison with the mouse lung.^16^ Mouse atlases cover aging, development, vascular and epithelial compartments, and influenza infection.^54–59^ Lung atlases have also been reported for pigs^60, 61^, dogs^62^, goats^63^, and 11 other non-model mammals, reptiles and birds^64^. In contrast, NHP lung atlases remain sparse, despite NHPs being an excellent model of human respiratory disease.^7, 8^ A single-nucleus atlas of young and aged cynomolgus macaque lungs annotated 19 cell types with T cells as a single population.^65^ A single-nucleus study of rhesus macaque lung, liver, kidney and brain after SARS-CoV-2 infection focused on broad cell classes.^20^ The Rhesus Immune Reference Atlas (RIRA) is limited to immune cells, with coarser annotation than current human lung references.^17^ The 2026 American Thoracic Society consensus report (C3LCN) now sets the standard for analysis, reporting and nomenclature of lung single-cell datasets, which it anchors to the healthy adult human lower respiratory tract.^66^ We followed this framework where applicable, although our annotation is somewhat less granular than the resolution it proposes. Our atlas of 37 cell types provides a highly detailed NHP lung reference, and the concordance of pigtail macaque and human lung populations supports its use as a translational model (Fig. S3). As mice appear to lack 17 of the 58 human lung cell types^16^, this concordance is relevant for studies in which rodent models may not capture human cellular composition.

The primary strength of this work is the detailed annotation of 37 cell types in the pigtail macaque lung, which provides a baseline for comparison with infected NHP models. Single-cell resolution is needed to study the cellular dynamics of respiratory pathogens such as influenza virus and SARS-CoV-2. These dynamics have mostly been examined in vitro^67^ or in mouse models^58, 59^, which differ from human in lung cell composition. Epithelial, B cell and monocyte populations in the pigtail lung corresponded closely to human and rhesus cell types. The atlas also resolves populations, which other references merge or do not label. RIRA assigned both IM and Mo-IM to alveolar macrophages (98% and 81%), and neither reference has separate labels for γδ T, MAIT-like and ILC3 populations. We used CellBender to remove technical artifacts and scVI for unsupervised hierarchical clustering, which may have contributed to the resolution of cytotoxic T, NK and innate lymphoid populations.^68^

Our cross-species mapping shows that neither the human nor the rhesus reference fully captures pigtail lung populations (Figs. S3 and S4). Weaker mappings were concentrated in macrophage, DC and innate-like T populations that the references merge or do not label. Human atlases share this limitation. Species differences in innate lymphocyte biology may contribute to lower correspondence to human innate lymphocytes. Macaque NK cells differ from human NK cells in CD56 and CD8α expression, which may explain why 41% of pigtail NK cells mapped to HLCA CD8 T cells (Fig. S4).^69^ The macaque KIR family has also expanded in a lineage-specific way and is not one-to-one orthologous with human KIR genes, which further limits ortholog-based mapping of NK subsets.^70^ However, macaques have phosphoantigen-reactive Vγ9Vδ2 T cells (like humans), which are early responders to respiratory viruses and not present in mice.^71–73^ Because mapping depends on the label sets and resolution of each reference, some differences may also reflect annotation granularity rather than divergent biology.

The main limitation is that this atlas was generated from five animals and only used lung parenchyma, which did not capture basal and goblet cells. The cohort also included only female macaques, so sex-related differences in lung cell populations remain incompletely characterized. A study modeling lung scRNA-Seq data determined that donor-specific effects were greater than sex-specific effects.^74^ Sex did not substantially affect cell-type classification performance in that study; however, sex differences may nevertheless emerge during infection. In human donors, females had significantly higher numbers of influenza-specific CD8+ terminally differentiated effector memory T cells that re-express CD45RA (TEMRA) and lower numbers of influenza-specific CD8+ TEM and tissue-resident memory cells than males.^75^

Beyond cohort composition, several technical limitations affect annotation of T and NK cell populations. Definitive annotation of MAIT cells in the pigtail macaque would require paired T cell receptor sequencing, CITE-seq or MR1 tetramer staining. Detection of genes and transcription factors that define CD4+ T cells and their subsets was poor, a known limitation of scRNA-seq datasets.^17^ Only a minority of CD4+ T cells contained CD4 transcripts, FOXP3 was detected in only a few CD4+Treg cells, and helper subsets (e.g., Th1, Th17) could not be distinguished. ILC1, ILC2 and CD16− NK populations were also not resolved, likely because of their low frequency in the adult lung and the number of cells captured. Finally, differences in data processing, clustering and depth of annotation between the NHP and human lung atlases limit direct comparison of resolution.

In summary, we describe a pigtail macaque lung atlas of 61,323 cells that resolves 37 cell types across epithelial, endothelial, stromal and immune compartments. Major lineages corresponded closely to human and rhesus reference atlases, with the highest concordance in alveolar epithelial, B cell and monocyte populations. The atlas also resolves macrophage and innate-like T populations that the references merge or do not label. It provides a reference for pigtail macaque studies of pandemic respiratory viruses and lung injury, including evaluation of vaccines and therapeutics.

## Methods

### Ethics Statement

All nonhuman primate experiments were performed in strict accordance with the recommendations in the Guide for the Care and Use of Laboratory Animals of the National Research Council and the Weatherall report: “The use of non-human primates in research”. The lung tissues were distributed through the Washington National Biomedical Research Center Tissue Distribution Program with approval from the Institutional Animal Care and Use Committee (IACUC) of the University of Washington (UW) under protocol 4277-01.

### Tissue Dissociation Pipeline

#### Single Cell Dissociation

Lung samples were immediately transferred into cold Hibernate-E (Thermo-Fisher, #A1247601) for transport and single-cell dissociation was performed within 1 hour after collection.

#### Isolation of Cells from the Lung

While submerged in a small volume of cold HBSS with calcium and magnesium (HBSS/Ca²⁺/Mg²⁺; Thermo-Fisher, #14025092), dissected tissues were finely chopped into approximately 2 to 3 mm pieces (∼100 to 1,000 mg) using sterile scissors and washed three times in HBSS/Ca²⁺/Mg²⁺ supplemented with Penicillin-Streptomycin (Fisher, #15-140-122) and Gentamicin/Amphotericin solution (Thermo-Fisher, #R01510). Digestion solution was freshly prepared using this same antibiotic-supplemented HBSS/Ca²⁺/Mg²⁺ base with 1:1000 diluted Collagenase IV stock (100 U/mL final; Fisher, NC9919937). Digestion was carried out at 37°C for 40–90 minutes in a water bath. Samples were vortexed every 15 minutes and digestion was considered complete once the solution appeared cloudy. Digestion was stopped with 10 mL of cold Wash/DNase solution, and samples were centrifuged at 1200 rpm for 3 minutes at room temperature.

Supernatant was aspirated, and the pellet resuspended in 3 mL of TrypLE™ Express Enzyme (Thermo-Fisher, # 12604013) and incubated in a 37°C water bath for 5 minutes. Subsequently, 10 mL of Wash/DNase solution was added, and samples were again centrifuged at 1200 rpm for 3 minutes at room temperature with the supernatant discarded. At this step, samples where a single tissue had been divided into multiple aliquots were pooled and washed three times in Wash buffer (2% heat-inactivated FBS in 1X PBS). The suspension was then passed through a cell strainer, centrifuged at 1200 rpm for 3 minutes at room temperature, and the pellet washed twice more with Wash buffer.

Red blood cells were lysed by resuspending the pellet in 2 mL of Ammonium Chloride solution (Fisher, NC9877880), followed by centrifugation at 1200 rpm for 3 minutes and two additional washes with Wash buffer. Cells were counted and viability was assessed prior to resuspension in the volume required for downstream application (for 10x Genomics scRNA-seq, a minimum of 80% viability and 700,000 fresh cells were required). Any remaining cells were cryopreserved in liquid nitrogen for long-term storage.

### Single Cell RNA-Seq Library Preparation

*Sequencing.* Single-cell RNA sequencing libraries were generated using the Chromium Next GEM Single Cell 3′ v3.1 Dual Index workflow (10x Genomics, Pleasanton, CA) following the manufacturer’s recommended protocol. Briefly, single-cell suspensions were combined with barcoded gel beads and reaction reagents on a Chromium Next GEM Chip G, and Gel Bead-In-Emulsions (GEMs) were produced using the Chromium Controller (10x Genomics, Pleasanton, CA). Cellular mRNA was reverse transcribed within each GEM. Then, cDNA was amplified and sequencing libraries were constructed according to the manufacturer’s guidelines. We assessed cDNA and library quality at multiple stages using an Agilent 2200 TapeStation system (Agilent Technologies, Santa Clara, CA). Libraries were initially subjected to shallow sequencing on a NextSeq 2000 platform (Illumina Inc., San Diego, CA) to assess alignment rates to the reference genome, GC content distribution, and sequencing depth on a per-cell basis. These metrics were used to determine whether libraries met quality requirements for deep sequencing and if sample balancing was needed to optimize read recovery per cell. Libraries that passed quality control were deep sequenced on a NovaSeq instrument (Illumina Inc., San Diego, CA), targeting approximately 30,000 reads per cell.

### Bioinformatics Pipeline

#### Alignment and preprocessing

Reads were aligned with Cell Ranger (v9.0.1) to the rhesus macaque Mmul10 genome with Ensembl release 113 protein-coding annotations.^76^ We used the rhesus reference because it is more complete than the pigtail macaque assembly and the two species are highly conserved in coding regions. Ambient RNA was removed with CellBender (v0.3.0).^68^ Cells were filtered based on detected genes, UMI counts, and mitochondrial fraction (calculated from the 13 mitochondrially encoded protein-coding genes, which lack an “MT-” prefix in Mmul10). Corrected matrices were loaded into Python (v3.10.19) using Scanpy (v1.11.5)^77^, and doublets were removed with Scrublet (v0.2.3)^78^.

#### Integration and clustering

Samples were integrated with scVI^79^ in scvi-tools (v1.3.3) using sample as the batch covariate, and the resulting model was refined with scANVI^80^ using preliminary Leiden clusters as labels. UMAP and Leiden clustering^81^ were run on the scANVI latent space. Integration improved sample mixing, with the median inverse Simpson index of sample identity among nearest neighbors rising from 1.40 to 2.27.

#### Cell type annotation

The integrated data were analyzed in Seurat (v5.5.0).^82^ Clusters were annotated using differentially expressed markers and canonical lineage markers, with heterogeneous clusters subclustered as needed. A second scANVI model trained on the annotated atlas was used to subcluster interstitial macrophages into Mo-IM, IM, and cDC2. UMAPs of each compartment were computed from this model’s latent space. Nomenclature followed the Collaborative Cross-Consortium and Country Lung Cell Nomenclature Project, an official initiative of the American Thoracic Society (ATS) Assembly Project.^66^

#### Cross-species comparison

Macaque cells were mapped to the HLCA core reference with scArches (v0.6.1)^83^ after conversion to human one-to-one orthologs, and labels were transferred using HCLA annotation level 4.^84^ scArches was used as the HLCA provides a pretrained reference model built for this purpose, which places new cells into the reference without retraining it. A transfer uncertainty was calculated for each cell and is reported as the median for each cell type.

Immune cells were also compared with the Rhesus Immune Reference Atlas (RIRA; GSE277821)^17^ by Seurat label transfer. RIRA does not provide a model of this kind, and because both datasets were aligned to the same rhesus genome, genes could be matched directly without ortholog conversion, which made anchor-based transfer the more direct choice. For this comparison, uncertainty was calculated as 1 minus the Seurat prediction score and is also reported as the median for each pigtail macaque cell type, with lower values indicating greater confidence in the transferred label.

## Declarations

### Availability of Data and Materials

The data generated and analyzed during the current study are available in the NCBI Gene Expression Omnibus (GSE348023). Bioinformatics code is available on the Adams Waldorf Lab GitHub (https://github.com/kadamswaldorflab/PTM_Lung_Atlas) or in the Dryad repository (DOI: 10.5061/dryad.r7sqv9svt). The lung atlas can also be explored using the UCSC Cell Browser: https://cells-test.gi.ucsc.edu/?ds=lung-atlas.

### Authors’ Contributions

KMAW supervised the research and analysis performed in this project, secured funding, analyzed the data, supervised the research team, and wrote and edited the manuscript. JAC also supervised aspects of this project. TK, JM, and JAC were involved in data curation, formal analysis, and software. AL was also involved in data curation. TK, RPK, and JAC performed the data analysis, created visual representations of the data, and interpreted the results. VS, TD, SS, SCC, HH, OC, HZ, BMDR, NB, EF, MW, DB, AL, AEV, AG, GM, AC, MRB, CE, AB, and JAC were investigators on this project and collected data. TK, VS, TD, SS, SCC, JAC, and KMAW wrote the first draft of the manuscript. All authors edited and revised subsequent drafts and approved the final manuscript.

## Acknowledgment

We thank the animal care, veterinary, and research support staff of the Washington National Biomedical Research Center for their expert care of the animals and their assistance with necropsy and tissue collection. We are grateful to the Washington National Biomedical Research Center Tissue Distribution Program for providing the lung tissues used in this study. We also acknowledge the nonhuman primates whose contributions made this research possible.

