## Supplemental Information for "A single-cell transcriptomic atlas of the pigtail macaque lung"

**Table S1.** Animal Demographics

| ID | Sex | Age (years) | Weight (kg) |
|---|---|---|---|
| NHP 1 | F | 8.8 | 11.2 |
| NHP 2 | F | 4.0 | 5.1 |
| NHP 3 | F | 3.8 | 5.1 |
| NHP 4 | F | 8.6 | 9.6 |
| NHP 5 | F | 11.2 | 10.4 |

This table shows the sex, age, and weight of the animal at the time of lung sampling and necropsy.

**Table S2.** Macaque-human gene symbol mapping for marker genes

| ENSMMUG ID | Assigned symbol | Human Ensembl ID | Notes |
|---|---|---|---|
| ENSMMUG00000003532 | *CD8B* | ENSG00000172116 | |
| ENSMMUG00000057791 | *TRDC* | ENSG00000211829 | |
| ENSMMUG00000063583 | *GNLY* | ENSG00000115523 | |
| ENSMMUG00000040771 | *IGHG3* | ENSG00000211897 | |
| ENSMMUG00000044861 | *IGLL5* | ENSG00000254709 | |
| ENSMMUG00000015202 | *IGHG1* | ENSG00000211896 | |
| ENSMMUG00000005378 | *TNFRSF13B* | ENSG00000240505 | |
| ENSMMUG00000061680 | *TRAV1-2* | ENSG00000256553 | Not detected; not shown in dot plots |

This table shows mapping between human and macaque genes used as markers for the lung atlas.

**Figure S1.** Quality control analyses of cell populations

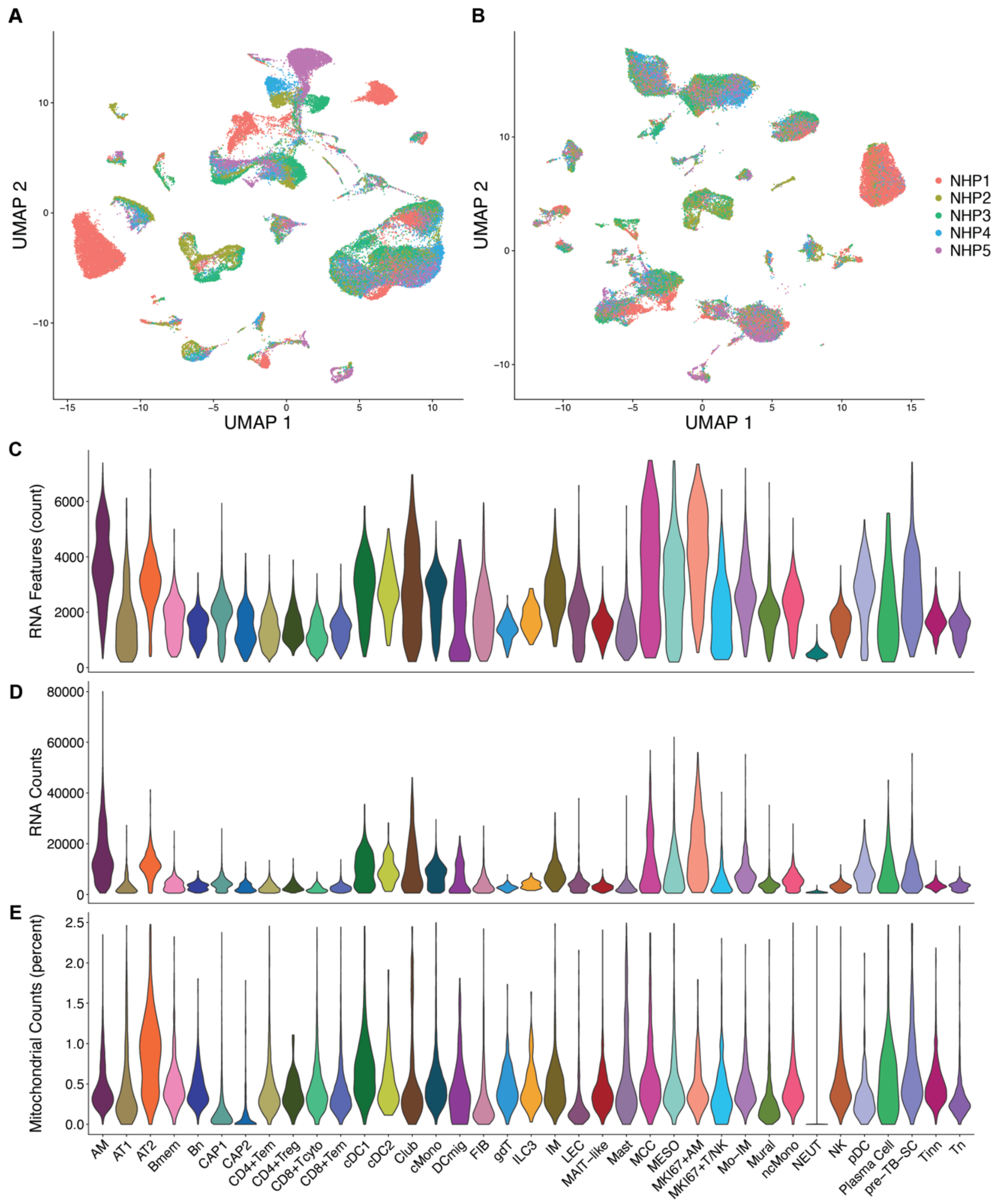

(A–B) UMAP embeddings of single-cell transcriptomes are colored by animal before (A) and after (B) scVI and scANVI integration. (C-E) Violin plots display the number of detected genes (C), number of unique molecular identifiers (UMI) counts (D), and percentage of mitochondrial counts (E) for each cell type. Cell type abbreviations are defined in the Figure 2 legend.

**Figure S2.** Cell Type Composition Plot Per Animal

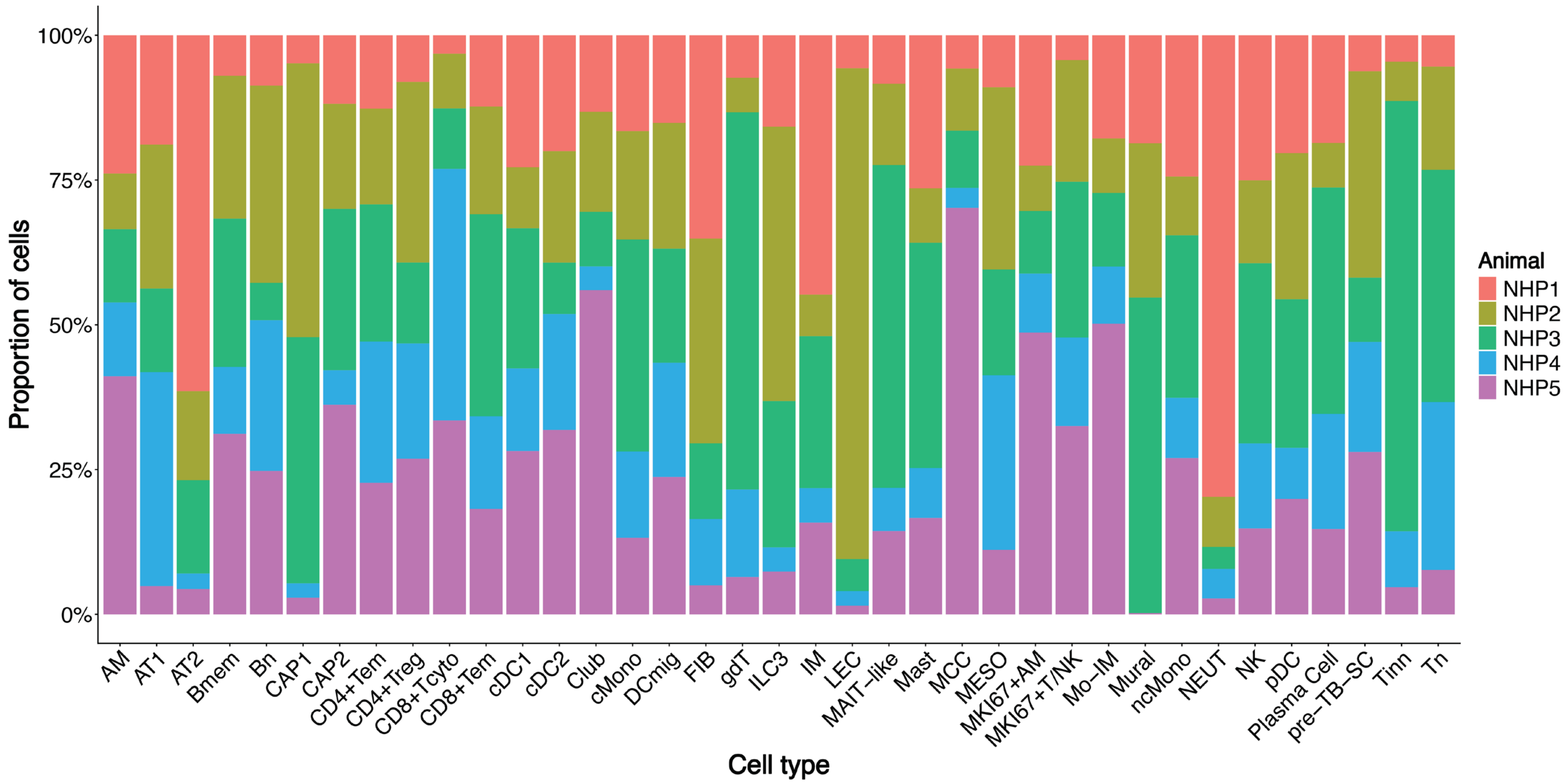

Stacked bar plot showing the proportion of cells contributed by each animal (NHP1 to NHP5) within each of the 37 annotated cell types. Each bar represents one cell type, and the colors denote individual animals. Several populations came mostly from a single animal, including lymphatic endothelial cells, neutrophils, innate-like CD8+ T cells, and multiciliated cells. Abbreviations for cell types are shown in Figure 2.

**Figure S3.** Cross-species mapping of the pigtail macaque lung atlas to the Level 4 Human Lung Cell Atlas

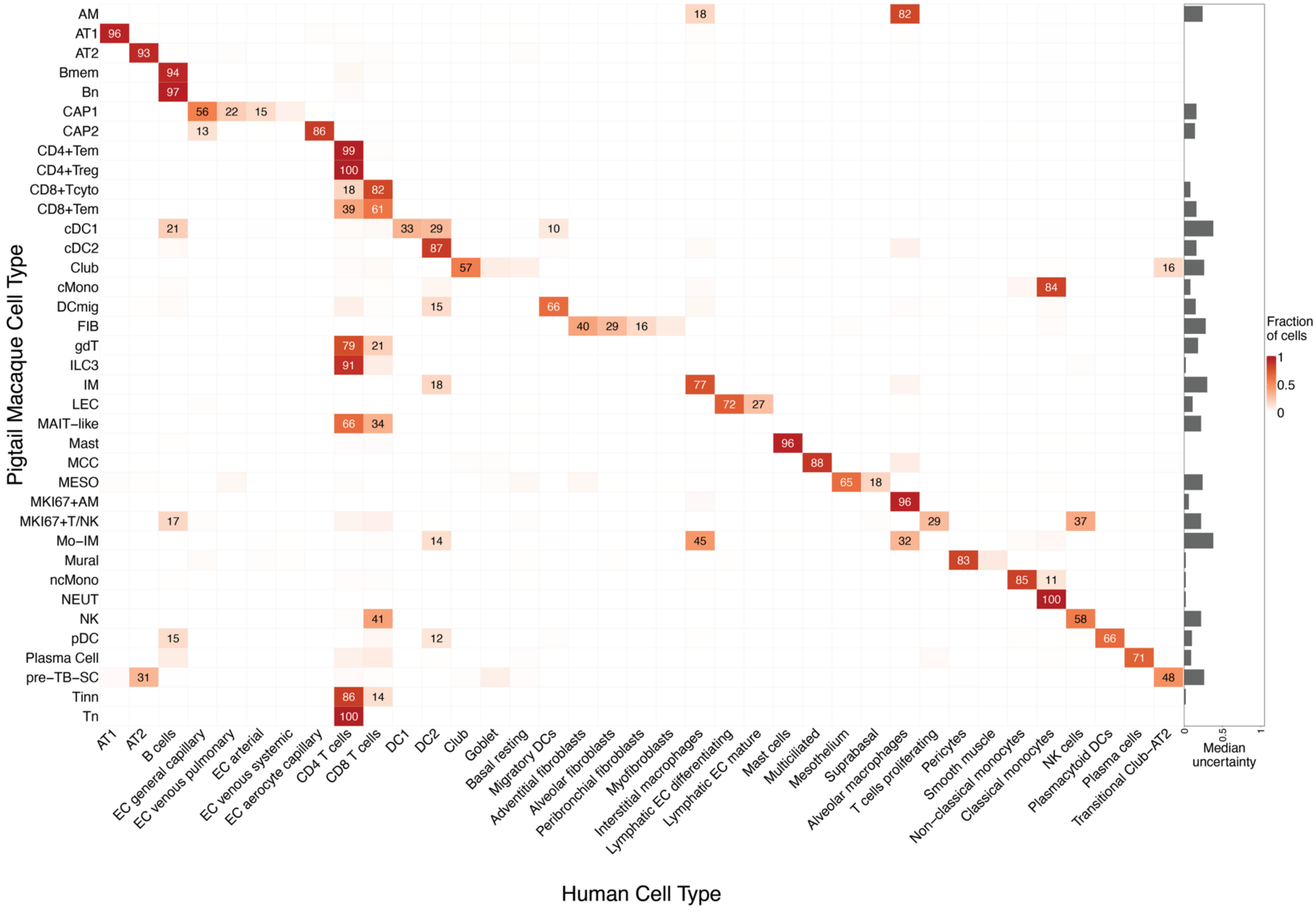

Pigtail macaque cell types on the y-axis were mapped to the Human Lung Cell Atlas (HLCA) level 4 using scArches. The estimated fraction of cells in each pigtail macaque cell type that would be assigned to each of the HLCA cell type is shown if the value is greater than 10% and assigned a color. The bar plot shows the median of 1 minus the prediction score for each pigtail macaque cell type, where lower values indicate greater confidence in the transferred label. Y-axis abbreviations are defined in the Figure 2 legend. Abbreviations: AT1, alveolar epithelial cell, type 1; AT2, alveolar epithelial cell, type 2; EC, endothelial cell; DC1, conventional type 1 dendritic cell; DC2, conventional type 2 dendritic cell; and NK, natural killer cell.

**Figure S4.** Cross-species mapping of pigtail macaque lung immune cells to Rhesus Immune Reference Atlas cell types

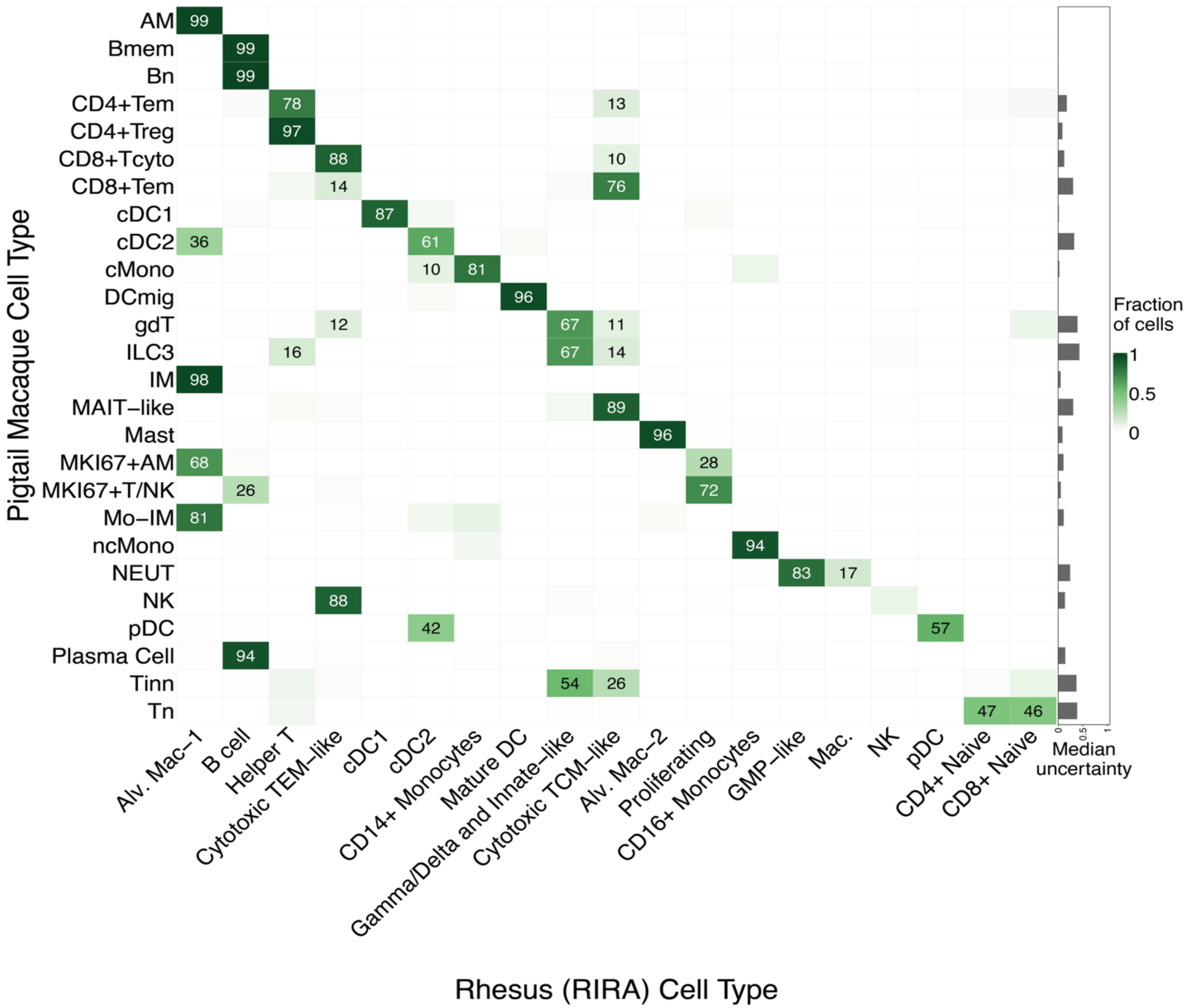

The y-axis labels are the 26 immune cell types annotated in the pigtail macaque lung atlas, and x-axis labels are Rhesus Immune Reference Atlas (RIRA) cell type annotations. The color gradient reflects the estimated fraction of cells in each pigtail macaque cell type assigned to each RIRA cell type. Values greater than 10% are labeled as percentages. The bar plot shows the median uncertainty (1 minus the prediction score) for each pigtail macaque cell type, where lower values indicate greater confidence in the transferred label. Row abbreviations are defined in the Figure 2 legend. RIRA abbreviations: Alv. Mac, alveolar macrophage; DC, dendritic cell; GMP, granulocyte-monocyte progenitor; Mac., macrophage; NK, natural killer cell; TCM, central memory T cell; TEM, effector memory T cell.
